# Novel monoclonal antibodies against the C-terminal HEAT domain of Huntingtin

**DOI:** 10.1101/2025.09.17.676860

**Authors:** Yang-Nim Park, Rebeka Fanti, Samira Sadeghi, Renu Chandrasekaran, Aled M. Edwards, Rachel J. Harding, Douglas W. Houston

## Abstract

**BACKGROUND:** Reliable detection of huntingtin (HTT) is essential for understanding Huntington’s disease (HD) biology and evaluating therapeutic strategies. However, high-quality monoclonal antibodies (mAbs) against the HTT C-terminal domain remain limited.

**OBJECTIVE:** We sought to generate and validate novel monoclonal antibodies targeting the HTT C-terminal HEAT-containing domain to better detect HTT independently of potential effects of polyglutamine length that can impact some N-terminally targeted antibodies.

**METHODS:** We immunized mice with a highly purified, well-characterized recombinant protein corresponding to the HTT C-terminal domain. We generated monoclonal antibody-producing hybridoma cell lines and characterized the antibodies using parental and HTT-knockout cell lines in common immuno-applications.

**RESULTS:** Three novel, independent hybridoma lines producing anti-HTT monoclonal antibodies were derived. Using CRISPR-edited HTT knockout cell lines we identified one clone, anti-HTT [2F8], that was specific and effective across Western blot, immunofluorescence, and ELISA assays. All antibodies bound full-length HTT irrespective of HAP40 interaction or polyQ length and showed no cross-reactivity to the N-terminal HEAT domain.

**CONCLUSIONS:** These C-terminal HTT mAbs are thus valuable additional tools for studying endogenous HTT function in both normal and disease contexts.

## Introduction

Current state-of-the-art therapies for Huntington’s disease (HD) aim to reduce levels of the mutant huntingtin protein (HTT), which contains an expanded polyglutamine (polyQ) tract in the exon one region of the protein.^1^ Accurately monitoring the efficacy of these approaches depends on assays that can reliably detect and quantify HTT protein in biofluids. Monoclonal antibodies (mAbs) are among the most widely used tools for protein detection, owing to their high specificity against discrete epitopes and their low cost and renewable nature.

HTT is a large, 348 kDa protein expressed ubiquitously across all tissues ^2^ and implicated in a wide range of intracellular processes, including vesicle recycling, endosomal trafficking, transcriptional and translational regulation, and coordination of cell division. ^3–5^ Despite broad functional associations, the precise molecular roles, direct binding partners, and sub cellular localization of different HTT-scaffolded complexes remain incompletely defined, making it challenging to understand how disease-causing mutations disrupt HTT activity and contribute to HD pathogenesis.

Structurally, HTT consists of a large N-terminal domain (NTD) and a smaller C-terminal domain (CTD), both of which contain HEAT repeats, connected by a central Bridge domain.^6^ HEAT repeats are structural motifs known for mediating dynamic protein–protein interactions due to their flexible architecture.^7^ Although numerous HTT interactors have been proposed, HAP40 remains the only structurally validated partner, and its expression levels follow the same patterns as HTT.^8^

Several mAbs targeting HTT are commercially available, however are directed against epitopes in the extreme N-terminus (N17), the NTD with various polyQ lengths, or the proline-rich Bridge domain.^9^ Consequently, there is a notable lack of well-validated antibodies targeting more downstream regions, particularly the CTD. This limitation hampers efforts to study the full-length and other proteoforms in a consistent fashion. Addressing this gap is essential to enable more comprehensive studies of HTT biology and to support the development of tools for both mechanistic insights and therapeutic monitoring.

## Materials and Methods

### Protein expression and purification

All expression constructs used here have been published before^6,10^ and are available through Addgene. HTT and HTT–HAP40 protein samples were produced in insect cells and purified following a protocol similar to that previously reported.^10^ In brief, Sf9 cells were infected with P3 recombinant baculovirus and cultured until cell viability decreased to 80–85%, typically around 72 hours post-infection. For preparation of the HTT–HAP40 complex, cells were coinfected at a 1:1 ratio with HTT and HAP40 P3 baculoviruses. Harvested cells were lysed by freeze–thaw cycles, and the lysate was clarified by centrifugation. HTT proteins were isolated using FLAG-affinity chromatography. The FLAG-eluted fractions were subsequently applied to a Heparin FF cartridge (GE), washed with 10 column volumes (CV) of 20 mM HEPES pH 7.4, 50 mM KCl, 1 mM TCEP, and 2.5% glycerol, and eluted over a gradient ranging from 50 mM to 1 M KCl across 10 CV. All samples underwent a final gel filtration step on a Superose 6 10/300 column equilibrated in 20 mM HEPES pH 7.4, 300 mM NaCl, 1 mM TCEP, and 2.5% (v/v) glycerol. For the HTT–HAP40 complex, an additional Ni-affinity purification step was included before gel filtration. Fractions corresponding to either the HTT monomer or the HTT–HAP40 heterodimer were pooled, concentrated, aliquoted, and flash-frozen for downstream experiments. Sample purity was evaluated by SDS–PAGE, and protein identities were verified by native mass spectrometry.

### Animals and immunizations

Mice were immunized as previously described.^11^ Briefly, two female BALB/c mice were immunized with purified recombinant HTT-CTD.^6^ Immunogen (30-50 µg) was combined with 50 µg of poly(I:C) (HMW VacciGrade, InvivoGen, San Diego, CA), 2 µg CpG oligos (5’-TCCATGACGTTCCTGACGTT-3’) (IDT, inc., USA), and 50 µg of anti-CD40 mAb (BioXCell, West Lebanon, NH) as the adjuvant,^12–13^ in 250 µl of sterile PBS. Mice were boosted three times at three weeks intervals and bled one week after each boost. Sera were tested at 1:500 dilutions for anti-HTT reactivity against C-terminal HTT in immunoblots. The mouse with the best anti-HTT reactivity was given a final boost three days before the spleen was harvested for hybridoma production. All procedures with mice were approved by the University of Iowa Institutional Animal Care and Use Committee (IACUC).

### Hybridomas

Hybridomas were generated as previously described.^11,13^ Briefly, splenocytes and myeloma fusion partner cells (Sp2/0-Ag14, ATCC) were each washed and strained and resuspended in serum-free RPMI 1640 (HyClone). Nucleated splenocytes and myeloma cells were mixed 5:1 and fused in 50% (v/v) PEG-RPMI 1640 medium at 37°C. Fused cells were washed in serum-free RPMI 1640 and dispersed into hybridoma culture medium [IMDM (HyClone), containing 15% fetal bovine serum (FBS), 20% RPMI 1640 medium conditioned by giant cell tumor (TIB-223; ATCC) and 1X HT-Hybri-Max (Sigma)]. Hybridoma cells were cultured overnight at 37°C before being divided into four 96-well plates for HAT selection. The culture medium, supplemented with 1x HAT (Sigma), was replaced every three days until the majority of wells were 25-50% confluent (about two weeks), when the supernatant was tested for IgG secretion.

Positive wells were expanded and tested for anti-HTT activity against the immunogen using immunoblots in primary screens. Monoclonal lines were obtained by limiting dilution to single cells and reactivity was confirmed by immunoblotting against C-terminal HTT protein and against wild type (WT) and HTT mutant cell lysates (HTT-KO). Antibody isotypes were determined from culture supernatants using the Mouse mAb Isotyping Test Kit (AbD Serotec), according to the manufacturer’s protocols.

### Cell lines and CRISPR genome editing

Wildtype and *HTT*-mutant HEK293T and DMS-53 cell lines were previously used to characterize anti-HTT antibodies in Fanti et al. 2024.^9^ An additional *HTT* mutant knockout (KO) cell line was generated for this study using U-87MG cells (ATCC product HTB-14, U-87MG ATCC; RRID:CVCL_0022). To create frameshifts and/or early stop codons in the *HTT* gene in U-87MG cells, a CRISPR-Cas9 guide RNA (5’-GACAGGAACGAGUAAGCCUG<u>UGG</u>-3’) targeting exon 8 was designed using online tools (Synthego; PAM is underlined, chr4: 3116082-3116101). Oligos comprising the guide RNA sequence were cloned into pSpCas9(BB)-2A-Puro (PX459) V2.0 (pSpCas9(BB)-2A-Puro (PX459) V2.0 was a gift from Feng Zhang (Addgene plasmid # 62988; RRID:Addgene_62988)).^14^ CRISPR/Cas9 mutagenesis was performed by transfecting U-87MG cells with the *HTT* CRISPR vector and followed by puromycin selection. A heterogeneous population was screened by sequencing the *HTT* exon 8 region, and the mutagenic spectrum assessed using ICE analysis (Synthego^15^). Clonal lines were derived by limiting dilution, resulting in the U-87MG ATCC *HTT*-KO c6 line 100% mutant for a 1 bp deletion (G), which causes a frameshift mutation and premature stop codon, p.297fsTer310 (numbering based on human HTT with 21 glutamines). This clonal line was used for anti-HTT antibody screening and validation. Cell lines were assessed for endogenous HTT expression using the Cancer Dependency Map (DepMap) portal DepMap.^16^

### Immunoblotting

Immunoblotting was performed by multiple methods, each previously described in Park et al. 2016^11^ and Ayoubi et al. 2024^17^. Briefly, cells were lysed in Cell Lysis Buffer or RIPA Buffer with DNA shearing manually (25-gauge needle) or sonication. Cell lysates (40-50 µg total protein) were run on premade Tris-Glycine-SDS gels (BioRad) or Bis-Tris gels with MOPS running buffer. Proteins were transferred to nitrocellulose, blocked in 5% non-fat dry milk in TBS/0.005% Tween-20 (TBS-T) and incubated overnight at 4°C in primary antibody diluted in blocking solution (0.5 µg/ml anti-HTT mouse and anti-GAPDH [2G7] rabbit monoclonal antibodies (DSHB; RRID:AB_2617426 and RRID:AB_3105905, respectively), 1:500 antiserum, or 1-10 – 1:100 test supernatants). After washing in TBS-T, blots were incubated in goat anti-mouse-IR Dye 800 and goat anti-rabbit-IR Dye s680 secondary antibodies (Licor, diluted 1:10,000 in blocking buffer) for one hour at room temperature (RT). After additional washing in TBS-T, signal was detected by direct scanning or by chemiluminescence (Licor Odyssey Fc).

### Immunofluorescence

For characterization of anti-HTT mAbs using U-87MG and DMS-53 cell lines, a mosaic WT/KO method standardized by the YCharOS initiative was used.^17^ Wildtype and KO cells were differentially labelled with green and red CellTracker dyes [CellTracker™ Green CMFDA (5-chloromethylfluorescein diacetate) and CellTracker™ Deep Red], respectively, mixed, and co-cultured for 2 days. Cells were then fixed in 4% PFA, washed with PBS, permeabilized with 0.1% Triton X-100 for 10 minutes at RT, and blocked with 10% goat serum in PBS + 0.1% Triton X-100. Cells were incubated overnight at 4°C with 2 µg/ml anti-HTT mAbs in PBS containing 5% goat serum and 0.01% Triton X-100, washed and incubated in Alexa 555-conjugated anti-mouse secondary antibodies (1:500, Invitrogen) for 1 hour at RT. Nuclei were counterstained with 1.0 µg/ml DAPI.

Images were acquired using a Leica TCS SP8 inverted confocal microscope incorporated into a Leica imaging system with LAS X software. Z-series stacks were collected at 1 µm intervals using a 20x objective. Fluorophores were excited using laser lines at 405 nm (DAPI), 488 nm (CMFDA), 552 nm (Alexa 555), and 638 nm (Deep Red). The obtained *\**.*lif* file z-series images were processed using FIJI software to generate z-projections and converted to TIFF format. Images for publication were prepared in Adobe Illustrator and Photoshop using only level and contrast adjustments. Other modifications included resizing, changing stroke/fill weights and colors, and annotated overlays.

### Immunoprecipitation

Immunoprecipitations were performed as described in.^11,17^ Cell lysates were prepared as for immunoblotting and cell debris was removed by centrifugation at 10,000 x g for eight minutes. Clarified lysates were diluted with lysis buffer to about 3.0 mg/ml and incubated with 5-8 µg of antibody pre-bound to protein A/G magnetic beads (500 µg; Bimake) for one hour at 4°C with rotation. Samples of starting material and unbound fractions were kept for later analysis. The immunoprecipitates were eluted in SDS-PAGE loading buffer by boiling, separated by SDS-PAGE, and analyzed by immunoblotting.

### ELISA

ELISA using recombinant HTT was performed as described in Harding et al. 2025.^18^ Briefly, recombinant proteins were diluted to 1 µg/mL using gel filtration buffer (20 mM HEPES pH 7.4, 300 mM NaCl, 2.5% glycerol, 1 mM TCEP at pH 7.4) and incubated in 96-well Nunc Maxisorp plates (Thermofisher Scientific, cat# 442,404) overnight at 4°C. Plates were washed 3 times with PBS-T with (1X PBS, 0.005% (v/v) Tween-20) and blocked with PBS-T containing 1% (w/v) BSA (blocking buffer) for 2 h at 37°C. Plates were incubated overnight at 4°C with 6-point 1:3 serial dilution of respective mAbs in triplicate. The plate was then washed 3 times with blocking buffer and incubated for 1 h at 37°C with HRP-conjugated goat anti-mouse IgG (H+L) secondary antibody (1:50,000; Invitrogen, cat# 31430). After washing 3 times with blocking buffer, 100 µL of 1X TMB substrate (Invitrogen, cat# N301) was added per well and incubated at RT for about 15 min. The reaction was then quenched with 100 µL of 1 M phosphoric acid. The absorbances were measured at 450 nm using the CLARIOstar microplate reader (BMG LABTECH). Data were fitted with the specific binding using the hill slope model in GraphPad Prism version 9.5.1.

## Results

### Generation of hybridomas producing anti-HTT C-terminal monoclonal antibodies

We used a previously characterized, highly purified recombinant protein comprising amino acids 2095-3138 of human HTT (HTT-CTD; **Fig. 1A**, mauve) as the immunogen to generate new anti-HTT mAbs. This protein is structurally identical to the corresponding region of full-length HTT protein.^6^ Two mice were immunized with HTT-CTD and both produced detectable immune response in the sera. Fusions were carried out for both mice, but only the fusion from the second mouse resulted in successful monoclonal hybridoma lines at the secondary screening stage. Overall, 2 x 10^8^ splenocytes were fused and 78 clones against HTT-CTD were generated during the initial screening stage.

**Fig. 1.**
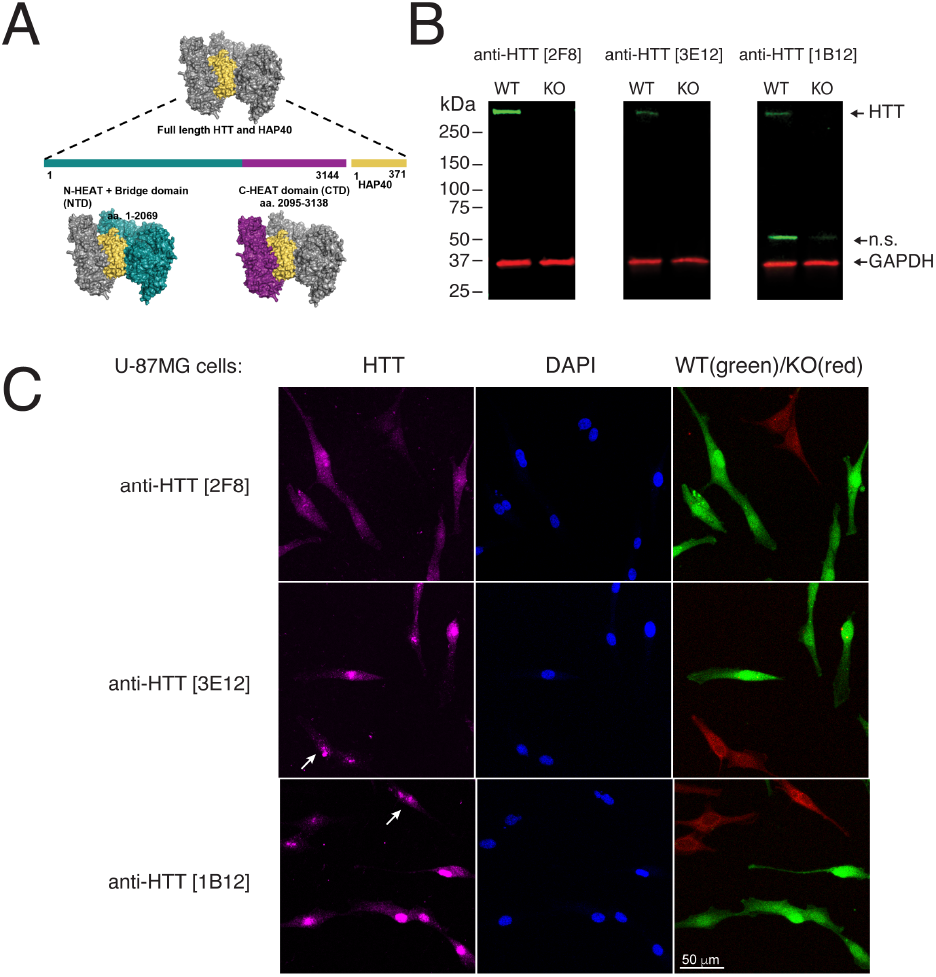
Characterization of anti-HTT monoclonal antibodies in U87MG cells. A) Diagram of human HTT in complex with HAP40. Colors indicate relative positions of domains and bound HAP40. B) Lysates of U87MG cells (wild type (WT) and HTT knockout (KO)) immunoblotted with the indicated antibodies. Anti-GAPDH [2G7] was used as a loading control. n.s.= non-specific band; relative migration is indicated (kDa). C) WT and KO U87MG cells labelled with green and far-red fluorescent dyes, respectively stained by IF using the indicated antibodies. Nuclei are indicated by counterstaining with DAPI. Arrows indicate non-specific nuclear signal in HTT-mutant cells. Scale bar = 50 µm.

We selected three hybridoma lines that showed robust reactivity against recombinant HTT-CTD protein and against endogenous HTT in U-87MG cells for further analysis: anti-HTT [2F8], [3E12], and [1B12]. Each of these lines secreted the IgG1 isotype and produced between 40-120 µg/ml antibody (**Table 1**). A fourth line was identified from a sibling subclone of anti-HTT [1B12], but analysis of immunoglobulin loci sequences later showed this clone was identical to 1B12.

**Table 1.** Summary of DSHB primary antibodies used.

| Antibody |  | Host Ig | Concentration |  |
| --- | --- | --- | --- | --- |
| anti-HTT [2F8] | AB_3712032 | mouse | 122 µg/mL | new |
| anti-HTT [3E12] | AB_3712033 | mouse | 88 µg/mL | new |
| anti-HTT [1B12] | AB_3712034 | mouse | 106 µg/mL | new |
| anti-HTT [2F8-Xi/Rabbit] | AB_3712035 | rabbit | 21 µg/mL | new |
| anti-HTT [3E12-Xi/Rabbit] | AB_3712036 | rabbit | 4 µg/mL | new |
| anti-GAPDH [2G7] | AB_2617426 | mouse | 62 µg/mL | new |
| anti-GAPDH [2G7-Xi/Rabbit] | AB_3105905 | rabbit | 34 µg/mL | new |

### Characterization of anti-HTT mAbs

Antibody characterization is most rigorous when incorporating gene knockout-based validation, the gold standard for ensuring specificity.^19^ To validate antibody specificity in U-87MG cells, we generated a clonal *HTT* knockout (*HTT*-KO) line by introducing a single nucleotide deletion in exon 8, leading to a frameshift and premature stop codon. Our initial immunoblots showed a predominant band at about 350 kDa, the size of full-length human HTT, in addition to lower MW bands. For all three mAbs tested, *HTT*-KO mutant U-87MG cells lacked the 350 kDa band (**Fig. 1B**). Lower MW bands, if present, were unchanged, as was the housekeeping protein GAPDH, demonstrating the specificity of the signal.

As a screen for specificity in additional assays, wildtype and knockout U-87MG cells were analyzed by IF. All three mAbs showed signal in both cytoplasm and nuclei of wild type U-87MG cells, although staining was almost exclusively nuclear for anti-HTT [3E12], and [1B12]. Anti-HTT [2F8] mAb lacked signal in mutant cells (**Fig. 1C**), whereas the others retained residual apparent nuclear signal and showed little difference with the control cells.

### Additional characterization of anti-HTT mAbs

To further assess the robustness of these mAbs across diverse contexts, we extended our analyses to additional human cell lines with differing origins and HTT expression profiles. This methodology can be beneficial because different levels of protein expression can lead to different results regarding the efficacy of the antibodies.^20^ HEK293 cells and DMS-53 cells were selected as fitting these criteria. HEK293 cells, although derived from kidney, display neuronal characteristics and express moderate levels of HTT,^21^ whereas DMS-53, a small cell lung carcinoma line, was selected based on an established dataset (DepMap) suggesting relatively high HTT abundance. In these lines, HTT was readily detectable in wild-type cells and absent in KO cells by immunoblotting. Similar to the U-87MG KO model, non-specific bands at 50 kDa were also observed (**Fig. 2A**). A summary of cell lines used is shown in **Table 2**.

**Table 2.** Summary of cell lines used.

| Cell line | RRID | Genotype | Reference | Source |
| --- | --- | --- | --- | --- |
| U-87MG<br>ATCC | CVCL_0022 | WT | - | ATCC |
| U-87MG<br>ATCC HTT<br>KO | CVCL_F0QD | KO | - | new |
| HEK293T | CVCL_0063 | WT | Ref. [9] | ATCC |
| HEK293T<br>_HTT null | CVCL_D7EP | KO | Ref. [9] | Academic |
| DMS-53 | CVCL_1177 | WT | Ref. [9] | ATCC |
| DMS-53<br>HTT KO | CVCL_D6U0 | KO | Ref. [9] | Academic |

**Fig. 2.**
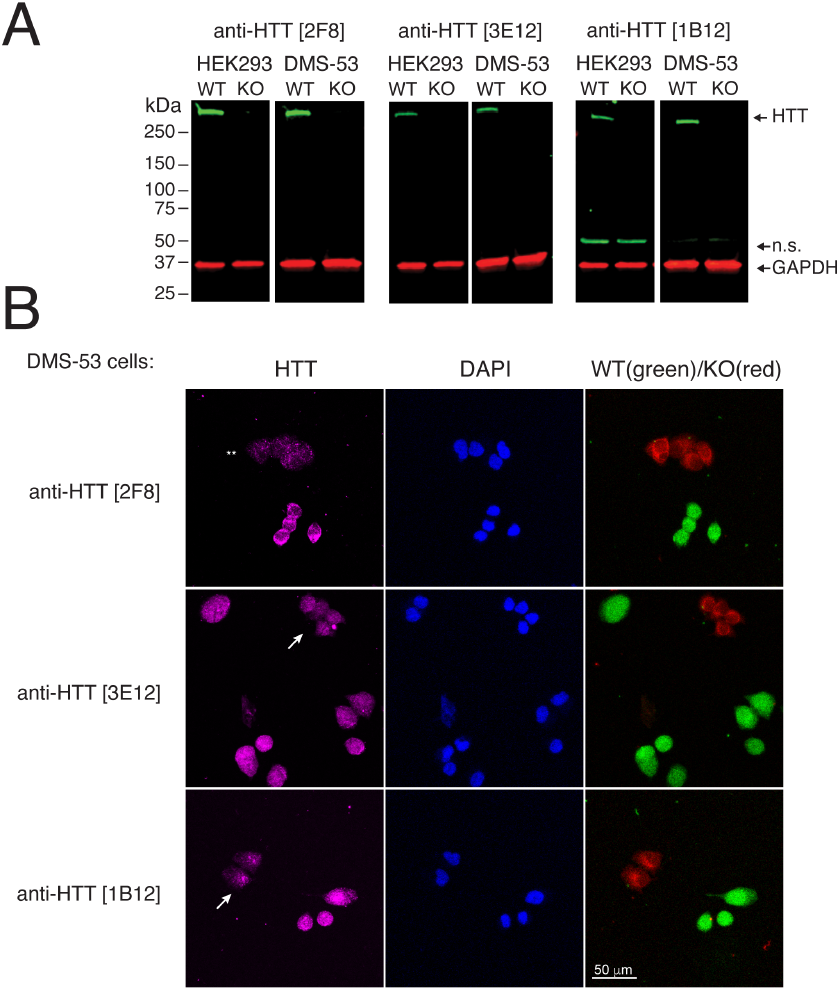
Characterization of anti-HTT monoclonal antibodies in additional cell lines. A) Lysates of HEK293T and DMS-53 cells (wild type (WT) and HTT knockout (KO)) immunoblotted with the indicated antibodies. Rabbit anti-GAPDH [2G7] was used as a loading control. n.s.= non-specific band; relative migration is indicated (kDa). B) WT and KO DMS-53 cells labelled with green and far-red fluorescent dyes, respectively stained by IF using the indicated antibodies. Nuclei are indicated by counterstaining with DAPI. Arrows indicate non-specific nuclear signal in HTT-mutant cells; asterisks indicate residual background staining for anti-HTT [2F8]. Scale bar = 50 µm.

Immunofluorescence was also used to assess antibody specificity and subcellular localization in additional cell lines. In wild-type DMS-53 cells, all three mAbs produced detectable signal in both the cytoplasm and nucleus. Similar to the case in U-87MG cells, non-specific nuclear signal was more pronounced for anti-HTT mAbs [3E12, and 1B12], and anti-HTT [2F8] clone demonstrated nearly complete loss of signal in the *HTT*-KO cells (**Fig. 2B**). The other clones retained residual nuclear staining or exhibited minimal differences between wild-type and KO cells, suggesting partial off-target binding. Background staining using anti-HTT [2F8] was slightly higher in the DMS-53 cells, but among the tested clones, anti-HTT [2F8] emerged as the best-performing mAb for detecting endogenous HTT across both immunoblot and IF platforms and across different cell lines.

To further expand the utility of these mAbs, we sequenced the immunoglobulin loci and cloned the variable regions of two clones (2F8 and 3E12) into rabbit Ig backbones. These recombinant chimeric (Xi/Rabbit) mAbs were tested in immunoblots using wild-type and KO cell lysates and were found to detect HTT in the same manner as the native mouse mAbs (**Fig. 3A**).

**Fig. 3.**
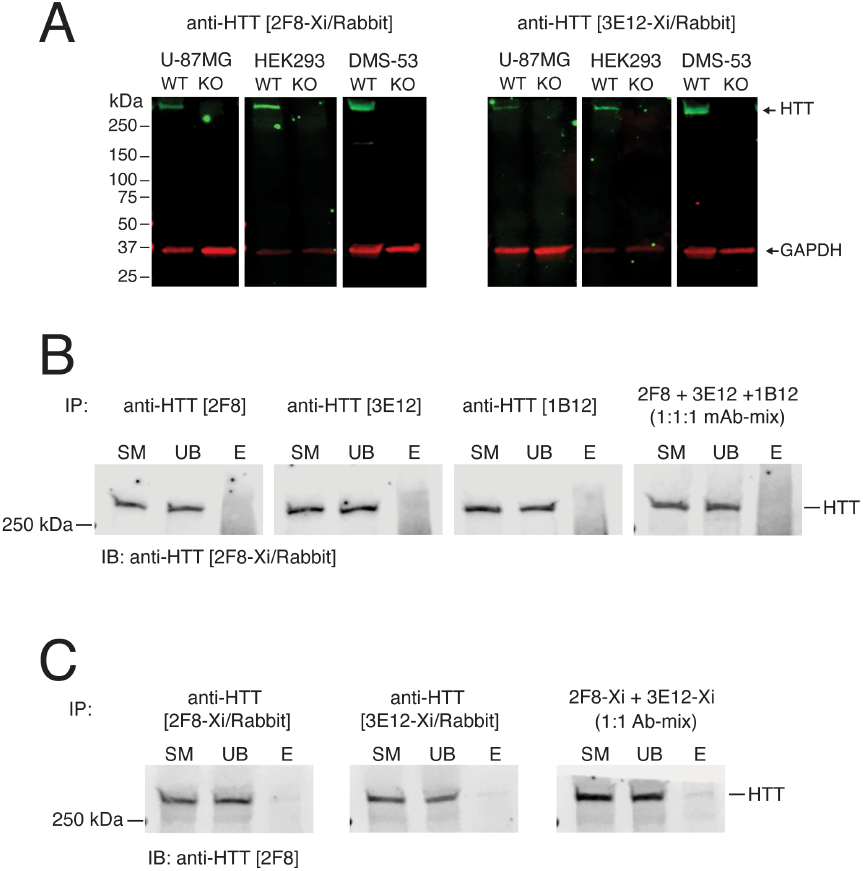
Characterization of chimeric rabbit anti-HTT monoclonal antibodies and screening by IP. A) Lysates of U-87MG, HEK293T, and DMS-53 cells (wild type (WT) and HTT knockout (KO)) immunoblotted with the indicated antibodies. Mouse anti-GAPDH [2G7] was used as a loading control; relative migration is indicated (kDa). B) Lysates of WT DMS-53 cells immunoprecipitated (IP) using the indicated mouse antibodies and immunoblotted (IB) using anti-HTT [2F8-Xi/Rabbit]. C) Lysates of WT DMS-53 cells immunoprecipitated (IP) using the indicated chimeric rabbit antibodies and immunoblotted (IB) using mouse anti-HTT [2F8]. SM, 5% starting material; UB, 5% unbound fraction; E, 100% eluted immunoprecipitate.

We next assessed whether anti-HTT mAbs were suitable for immunoprecipitation (IP) of endogenous HTT. Clarified lysates from wild-type DMS-53 cells were incubated with each mouse antibody, and the resulting immune complexes were analyzed by immunoblotting using an orthogonal rabbit antibody (anti-HTT [2F8-Xi/Rabbit] (**Fig. 3B**). The mouse mAbs only pulled down a diffuse smear of protein and failed to pull down detectable levels of HTT or visibly deplete the protein from the sample, even when used in combination (**Fig. 3B, right panel**). In the context of IP, recovery of >10% is typically considered efficient, particularly for large, multi-domain proteins such as HTT. The rabbit chimeric mAbs produced slightly cleaner IPs, with a weak band being detected in each case. However, the unbound supernatant following IP remained undepleted (**Fig. 3C**). Thus, these mAbs are not considered effective for IP, at least for endogenous full-length HTT in cell lysates.

Overall, these results demonstrate that monoclonal antibodies targeting the C-terminal domain of HTT can be effectively generated and validated using gene-edited knockout controls. The antibodies show utility across several key applications— including immunoblotting and immunofluorescence and are compatible with different cell line models.

### Anti-HTT mAbs Recognize HTT Independently of HAP40-bound state and PolyQ Length

Interactions between the N- and C-terminal HEAT domains of HTT and the 40-kDa huntingtin-associated protein (HAP40; *alias* F8A1 (coagulation factor VIII associated 1)) stabilize the compact and globular conformation of HTT.^8^ Because this interaction might shield the epitope recognized by our mAbs, which were raised against apo HTT-CTD, we evaluated the extent the C-terminal mAbs would recognize both HAP40-bound and apo forms of HTT, using a previously established ELISA-based binding assay.^18^

A panel of recombinant proteins, including recombinant CTD, CTD in complex with HAP40 (CTD-HAP40) and full-length HTT variants either with non-expanded (Q23-HAP40) or with expanded polyQ (Q54-HAP40) were tested for cross-reactivity with all three mAbs (**Fig. 4**). We included the N-terminal domain (NTD) of HTT as a negative control. All three antibodies robustly recognized CTD, CTD-HAP40, and full-length HTT-HAP40 complexes, regardless of polyQ length. Binding occurred in a dose-dependent manner, consistent with high-affinity and saturable antigen recognition. Importantly, no mAbs bound the NTD, confirming their specificity for epitopes located in the C-terminal region of HTT (**Fig. 4**). Further, clone anti-HTT [2F8] exhibited highest affinity for HTT and/or HTT-HAP40 complex. These results suggest that the selected mAbs recognize epitopes preserved in both the free and HAP40-bound conformations of HTT, which is recognized independently of the polyQ tract.

**Fig. 4.**
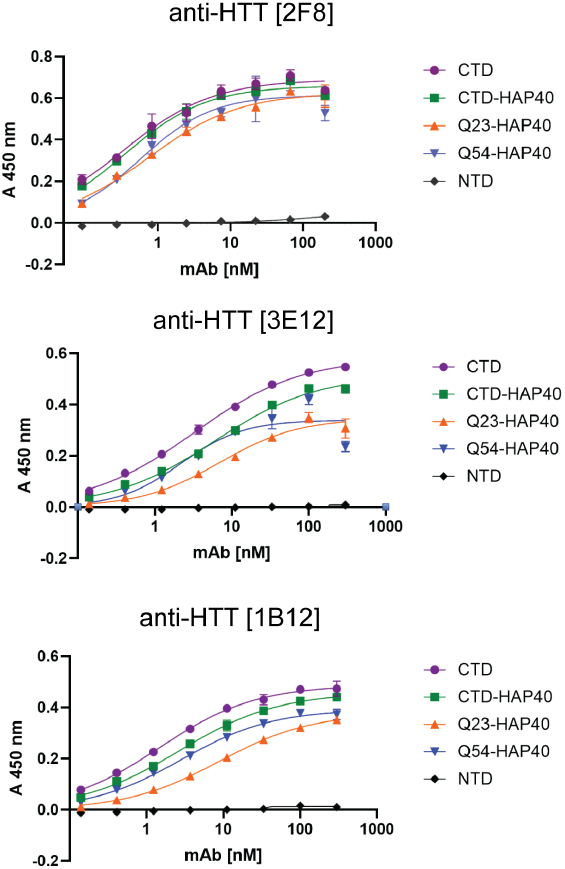
ELISA-based binding assessment using recombinant HTT proteins differing in domain composition, polyQ length, and the presence or absence of HAP40. The indicated proteins were diluted to 1.0 µg/mL and incubated with serial dilutions of anti-HTT mAbs. The plots show TMB-based detection measured at 450 nm. Values represent means of three replicates; error bars indicate standard deviations.

## Discussion

We have developed three novel antibodies against the C-terminal domain of human huntingtin, which we assessed in the gold standard knockout validation pipeline. We anticipate that these reagents will provide more high-quality tools reliable detection and quantification of HTT independently of changes in polyQ length and association with other proteins. The hybridomas we have developed are being deposited with the Developmental Studies Hybridioma Bank, a resource dedicated to the open-science sharing of monoclonal antibodies for research (see Ref. [22]). Plasmids will also be shared through Addgene, and we hope that these “open source” antibodies^23^ will help accelerate research on treatments for Huntington’s disease.

In developing these monoclonal antibodies, we took inspiration from the efforts of the YCharOS consortium (in which the authors’ organizations also participate), which aims to perform an open-science characterization of mAbs in popular applications by testing wild-type and mutant cells in parallel.^20^ We incorporated the knockout characterization steps into the initial hybridoma screening procedure so that only mAbs with a high probability of being specific for the antigen would be pursued. We identified three hybridoma clones that produced mAbs capable of specifically detecting endogenous HTT in three cell lines, each of a different lineage and level of HTT expression.

Even so, further testing of these mAbs in immunofluorescence assays showed that two did not show suitable specificity in two different knockout cell lines, and none of the antibodies effectively immunoprecipitated HTT from cell lysates. One clone, anti-HTT [2F8], was effective across immunoblot, immunofluorescence and ELISA assays.

It is possible that some of these antibodies will be effective in other assays, and deeper screening of anti-HTT hybridoma clones may be needed to identify specific mAbs to cover a more complete range of assays. Future experiments will help determine whether anti-HTT [2F8] works against HTT in paraffin or frozen sections or in applications like flow cytometry. To date, we have not observed reaction of any of these mAbs with mouse Htt. These antibodies may thus be human-(or primate-) specific. Importantly, however, all antibodies described here bind to full-length HTT irrespective of HAP40 interaction or polyQ length, and do not cross-react to the N-terminal HEAT domain of HTT in ELISA assays. These antibodies are thus expected to be useful for studying endogenous HTT function in both normal and disease contexts.

Another strategy we found productive was to use recombinant protein produced for the Structural Genomics Consortium (SGC) as our immunogen. The SGC is another open-science initiative dedicated to producing open-source 3D structures of potential drug target proteins. Because these proteins are highly purified and have been structurally characterized as part of SGC, they make ideal antigens for deriving antibodies active against native proteins. We expect to further develop this strategy with other proteins studied as part of the SGC, in particular with respect to understudied proteins implicated in human neurological or other diseases.

## Acknowledgments

The authors would like to acknowledge the YCharOS Consortium for developing and inspiring the knockout validation approach. Also, we thank M. E. MacDonald and I. S. Seong from Massachusetts General Hospital for a gift of *HTT*-null HEK293T cells. The following community resources supported this work: Addgene (RRID:SCR_002037), DSHB (RRID:SCR_013527), and the Structural Genomics Consortium (RRID:SCR_003890).

## Author contributions

Y-N, P. generated the hybridomas, performed and interpreted experiments and drafted the manuscript; R.F. performed and interpreted experiments and drafted the manuscript; S.S performed and interpreted experiments, R.C. performed and interpreted experiments, A.M.E. obtained funding and interpreted experiments, R.J.H. obtained funding, interpreted experiments, wrote the final manuscript draft; D.W.H. conceived the study, obtained funding, interpreted experiments, wrote the final manuscript draft.

## Statements and declarations

### Ethical considerations

All animal protocols used in these studies were approved by the Institutional Animal Care and Use Committee (IACUC) at The University of Iowa. PHS Animal Welfare Assurance (D16-00009, A3021-01).

